# Graph lesion-deficit mapping of fluid intelligence

**DOI:** 10.1101/2022.07.28.501722

**Authors:** Lisa Cipolotti, James K Ruffle, Joe Mole, Tianbo Xu, Harpreet Hyare, Tim Shallice, Edgar Chan, Parashkev Nachev

**Author notes:** Correspondence to: Prof. Lisa Cipolotti, Department of Neuropsychology, National Hospital for Neurology and Neurosurgery, Queen Square, London. WC1 3BG. Abbreviations: AH4-1 = Part 1 of the Alice Heim 4; APM = Advanced Progressive Matrices; Gf = Fluid intelligence; GNT = Graded Naming Test; LF = left frontals; LNF = left non-frontals; MD = multiple-demand network; NART = National Adult Reading Test; P-FIT = parieto-frontal integration theory; PLSM = parcel-based lesion symptom mapping; RF = right frontals; RNF = right non-frontals; ROI = regions of interest; WAIC = widely applicable information criterion; WAIS = Wechsler Adult Intelligence Scale.

## Abstract

Fluid intelligence is arguably the defining feature of human cognition. Yet the nature of its relationship with the brain remains a contentious topic. Influential proposals drawing primarily on functional imaging data have implicated “multiple demand” frontoparietal and more widely distributed cortical networks, but extant lesion-deficit studies with greater causal power are almost all small, methodologically constrained, and inconclusive. The task demands large samples of patients, comprehensive investigation of performance, fine-grained anatomical mapping, and robust lesion-deficit inference, yet to be brought to bear on it.

We assessed 165 healthy controls and 227 frontal or non-frontal patients with unilateral brain lesions on the best-established test of fluid intelligence, Raven’s Advanced Progressive Matrices, employing an array of lesion-deficit inferential models responsive to the potentially distributed nature of fluid intelligence. Non-parametric Bayesian stochastic block models were used to reveal the community structure of lesion deficit networks, disentangling functional from confounding pathological distributed effects.

Impaired performance was confined to patients with frontal lesions (*F*(2,387) = 18.491; *p* < .001; frontal worse than non-frontal and healthy participants *p* < .01; *p* <.001), more marked on the right than left (*F*(4,385) = 12.237; *p* < .001; right worse than left and healthy participants *p*<.01; *p*<.001). Patients with non-frontal lesions were indistinguishable from controls and showed no modulation by laterality. Neither the presence nor the extent of multiple demand network involvement affected performance. Both conventional network-based statistics and non-parametric Bayesian stochastic block modelling heavily implicated the right frontal lobe. Crucially, this localisation was confirmed on explicitly disentangling functional from pathology-driven effects within a layered stochastic block model, prominently highlighting a right frontal network involving middle and inferior frontal gyrus, pre- and post-central gyri, with a weak contribution from right superior parietal lobule. Similar results were obtained with standard lesion-deficit analyses.

Our study represents the first large-scale investigation of the distributed neural substrates of fluid intelligence in the focally injured brain. Combining novel graph-based lesion-deficit mapping with detailed investigation of cognitive performance in a large sample of patients provides crucial information about the neural basis of intelligence. Our findings indicate that a set of predominantly right frontal regions, rather than a more widely distributed network, is critical to the high-level functions involved in fluid intelligence. Further they suggest that Raven’s Advanced Progressive Matrices is a useful clinical index of fluid intelligence and a sensitive marker of right frontal lobe dysfunction.

## Introduction

Fluid intelligence (Gf) refers to the ability to solve challenging novel problems when prior learning or accumulated experience are of limited use.^1^ Gf ranks amongst the most important features of cognition, correlates with many cognitive abilities (e.g. memory),^2^ and predicts educational and professional success,^3^ social mobility,^4^ health^5^ and longevity.^6^ It is thought to be a key mental capacity involved in ‘active thinking’, ^7^ Gf declines dramatically in various types of dementia^8^ and reflects the degree of executive impairment in older patients with frontal involvement.^9^ Despite the importance of Gf in defining human behaviour, it remains contentious whether this is a single or a cluster of cognitive abilities and the nature of its relationship with the brain.^10^

Gf is traditionally measured with tests of novel problem-solving with non-verbal material that minimize dependence on prior knowledge. Such tests are known to have strong Gf correlations in large-scale factor analyses.^11, 14^ Raven’s Advanced Progressive Matrices^12^ (APM), a test widely adopted in clinical practice and research,^13^ contains multiple choice visual analogy problems of increasing difficulty. Each problem presents an incomplete matrix of geometric figures with a multiple choice of options for the missing figure. Less commonly, verbal tests of Gf such as Part 1 of the Alice Heim 4 (AH4-1)^15^ are adopted. The Wechsler Adult Intelligence Scale (WAIS) ^16^ has also been used to estimate Gf by averaging performance on a diverse range of sub-tests. However, several sub-tests (e.g. Vocabulary) emphasize knowledge, disproportionately weighting measures of “crystallized” intelligence,^17, 18^ whilst others (e.g. Picture Completion) have rather low Gf correlations.^19^ Hence, it has been argued that tests such as the APM are the most suitable for a theoretically-based investigation of changes in Gf after brain injury.^20, 21^

Proposals regarding the neural substrates of Gf have suggested close links with frontal and parietal functions. For example, Duncan and colleagues^22^ have argued that a network of mainly frontal and parietal areas, termed the ‘multiple-demand network’ (MD), is “the seat” of Gf. The highly influential parieto-frontal integration theory (P-FIT), based largely on neuroimaging studies of healthy subjects, posits that structural symbolism and abstraction emerge from sensory inputs to parietal cortex, with hypothesis generation and problem solving arising from interactions with frontal cortex. Once the best solution is identified, the anterior cingulate is engaged in response selection and inhibition of alternatives^23, 24^. Despite its name, P-FIT also posits occipital and temporal involvement, implying widely distributed substrates of Gf.^25^ A modification to P-FIT proposes a closer connection between frontal than parietal, regions and Gf-related processes,^26, 27^ with the frontal lobes mediating high-Gf “domain-independent” executive processes whilst posterior areas, including the parietal lobes, mediating low-Gf “domain-dependent” processing of spatial, object, or verbal information.

A meta-analysis of the functional imaging literature has implicated a network of modality-independent regions involving the inferior and middle frontal and inferior parietal lobes, with additional frontal eye field activation in non-verbal tasks, and anterior cingulate and left inferior frontal activation in verbal tasks.^28^ This fronto-parietal attention network,^29^ requires expansion to account for the separate neuronal substrate underpinning visuospatial/verbal analytical reasoning^30, 31^.

An important caveat of the functional imaging findings is that they do not imply causal efficacy.^32^ For example, though neuropsychological data commonly lateralise language to the left hemisphere,^33^ neuroimaging activation is often bilateral.^34^ So, merely considering the presence or absence of activation may hide lateralized functions. Hence, lesion studies offer an advantage in furthering our understanding of the neurocognitive architecture underpinning Gf. So far, these studies have been surprisingly sparse.

Lesion studies investigating performance on Gf tasks have mainly enrolled veterans with penetrating head injury.^35–43^ For example, Weinstein and Teuber ^43^ reported that veterans with left temporo-parietal entrance wounds suffered a decline in Army General Classification Test scores. Barbey and colleagues^44^ investigated Veterans’ performance on three subtests from the WAIS (Matrix Reasoning, Block Design and Picture Completion). The authors reported that performance was associated with damage to the superior longitudinal/arcuate fasciculus. However, several of the tests adopted are not considered Gf measures^45^; the lesion characterisation was rather basic and lacked modelling of the diffuse axonal injury the traumatic aetiology implies.^46^ In the most recent of these studies, impaired WAIS’s performance, not a specific test of Gf, was associated with damage to left fronto-parietal regions and white matter association tracts lesions.^47^ Glascher and colleagues^49^ reported that the left frontal pole was associated with performance on general intelligence (g) in a large sample of patients with stroke, encephalitis, temporal lobectomy and TBI. Similarly, Browen and colleagues^50^ investigated performance in general intelligence using the WAIS in a sample of patients with similar pathologies, and reported an association with white matter tracts deep to the left temporo-parietal junction, including the arcuate fasciculus.

Studies investigating patients with lesions caused by brain tumours or stroke have generally relied on WAIS as a measure of Gf, with inconclusive results. Some studies have associated WAIS non-verbal scale performance with right posterior damage.^48, 51^ However, Tranel and colleagues^52^ reported no significant differences between frontal and non-frontal damage on a non-verbal subtest of the WAIS analogous to the APM (Matrix Reasoning). Preserved performance on the WAIS has been documented in frontal patients.^53, 54^ In contrast, the very few studies adopting tasks loading more heavily on Gf have reported frontal deficits. Duncan and colleagues^20^ documented a substantial discrepancy between scores on Scale 2 of Cattell’s Culture Fair and the WAIS in 3 frontal but not in 5 non-frontal patients. However, the very small sample limited generalizability and prevented investigation of laterality effects.

In a recent study we documented lateralised frontal effects on APM and AH4-1.^55^ Compared with healthy participants, only right frontal damage significantly impaired APM performance, and only left frontal damage impaired AH4-1 performance. The relatively small sample prevented investigation of finer anatomical effects.

Lesion studies investigating the underlying behavioural and anatomical aspects of the widely used APM are old, inconclusive, and lacking in anatomical analysis. Results have variously shown no difference between right or left hemisphere patients;^56–61^ or impairment in left hemisphere patients^56, 62, 63^ or right hemisphere patients.^56, 57, 64–65^. Villardita^67^ reported no difference in the performance of left or right hemisphere patients on the Coloured Progressive Matrices version of APM. However, on set I, involving visuoperceptual factors, right performed worse than left hemisphere patients. There is similar uncertainty about the influence of aphasia, thought by some to degrade performance,^58, 68^ but not by others.^60, 62, 63, 69, 70^. Large samples of patients with focal unilateral lesions, thorough investigation of performance on the APM, fine-grained anatomical mapping, and robust lesion-deficit inference are vital for definitive scientific conclusions.

Here we assessed the largest number of patients yet reported with focal, unilateral, right or left, frontal or non-frontal lesions (n=227; 146 Frontal, 81 Non-Frontal, 165 healthy participants) on APM. We investigated overall performance, item difficulty, and relation to MD involvement. Building upon our novel multimodal methodology,^71^ we employed an array of lesion-deficit models responsive to the potentially distributed nature of Gf. We focused on modelling the anatomy of neural dependence as a graph, where interactions between distributed areas are explicitly tested. This approach permits delineation of distributed substrates. It also distinguishes functionally critical areas from those the distinctive pathological structure of lesions renders spuriously correlated: a problem shown to corrupt lesion-deficit maps based on simple mass univariate methods.^72^ In our approach each brain locus— intact or lesioned—is conceived as a node or vertex of a graph, with the relationships between loci— functional or merely lesion-pathology driven—defining its edges. This permits us to model network-dependence explicitly, disentangling functional and pathological effects to reveal the underlying substrate.

## Materials and methods

### Participants

Data from 332 patients with unilateral, focal lesions who attended the Neuropsychology Department of the National Hospital for Neurology and Neurosurgery was retrospectively screened. Inclusion criteria were: presence of a stroke or tumour; ≥70% of the total lesion, segmented from MRI or CT scans obtained during routine clinical care (see neuroimaging investigations), falling within either frontal or non-frontal areas; age between 18-80 years; absence of gross perceptual impairments (no neglect, >5^th^ cut-off on the Incomplete Letters test),^73^ language impairments (>5^th^ %ile on the Graded Naming Test, GNT);^74^ psychiatric disorders, history of alcohol or substance abuse, or other neurological disorders; and native English language proficiency. Age at assessment, gender and years of education were recoded.

Application of these criteria yielded 227 patients, 146 Frontal (LF 69; RF 77), and 81 Non-Frontal (LNF 39; RNF 42; see Table 1). There was no significant difference between tumour and stroke patients for mean time between injury and neuropsychological assessment (*p* = 0.12; table 1). 165 healthy control participants, with no neurological or psychiatric history, were recruited to match patients as closely as possible for age, gender, years of education and National Adult Reading Test scores (NART) ^75^.

**Table 1.**
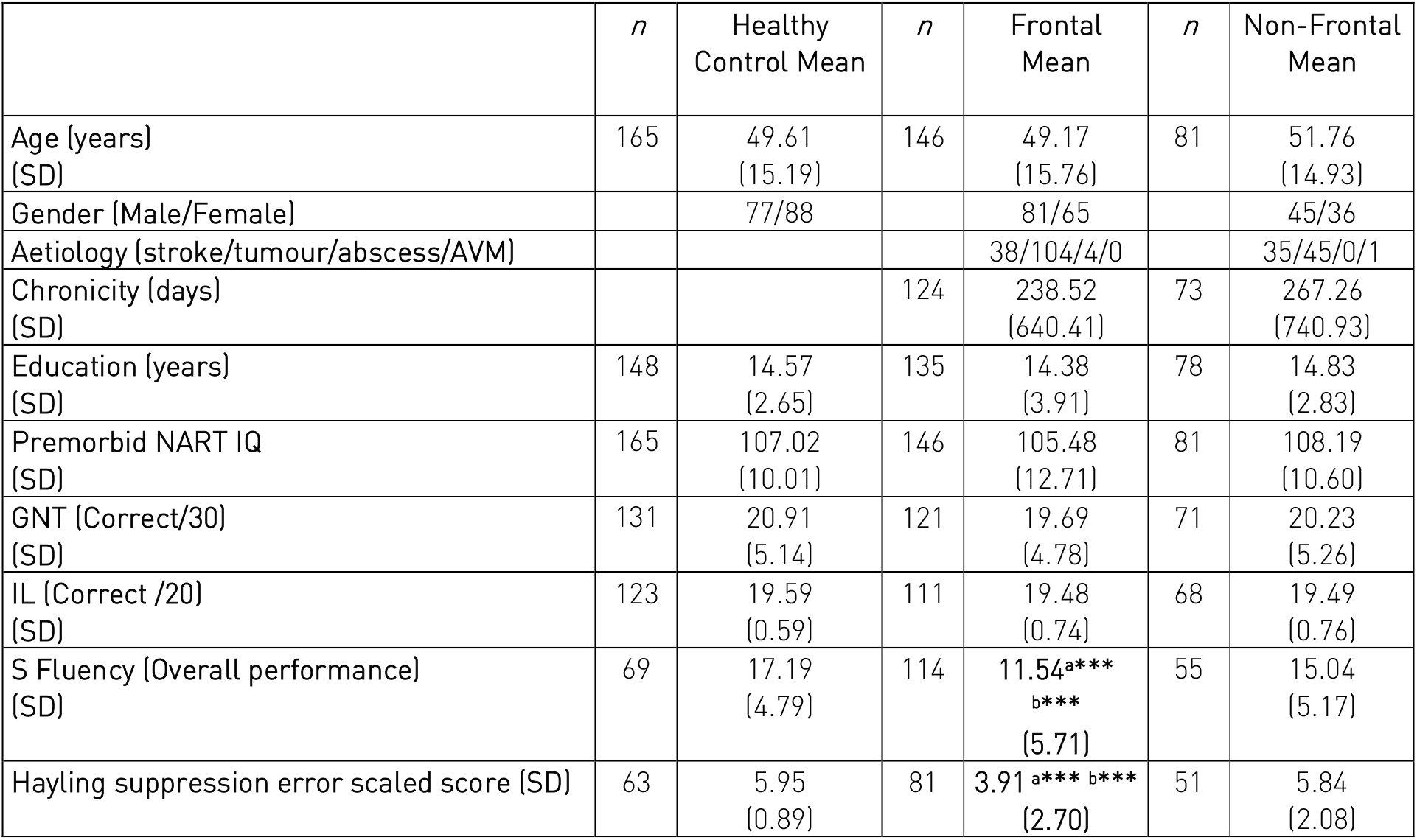
Demographics and cognitive test scores. n = Number. SD = Standard Deviation. AVM = arteriovenous malformation. NART= National Adult Reading Test. GNT = Graded Naming Test. IL=Incomplete letters. Scores with significant p values are in bold. ***= p <0.001. ^a^ indicates significant difference between frontal and non-frontal patients. ^b^ indicates significant difference between frontal and healthy control patients.

The study was approved by The National Hospital for Neurology and Neurosurgery & Institute of Neurology Joint Research Ethics Committee and conducted in accordance with the “Declaration of Helsinki”.

### Behavioural investigations

Patients were assessed with tests administered and scored in the published standard manner. Due to the retrospective nature of our study, certain data were unavailable for some participants.

#### Background tests

Premorbid optimal level of functioning was assessed using the NART, perception and naming using Incomplete Letters and GNT. Two widely used executive tasks, known to require processes distinct from Gf were also administered.^9, 76^ Verbal generation was assessed using the phonemic fluency test.^77^ The total number of words recalled excluding errors was recorded. Strategy formation/response inhibition was assessed using the Hayling Sentence Completion Test. Suppression Errors in Section 2 were calculated.^78^

#### Fluid intelligence

Gf was assessed using APM.^12^ We analysed the following:

a. *Overall performance.* The total number of correct responses in Set 1 (/12) was recorded and converted into age-adjusted scaled scores based on published norms.
b. *Item difficulty.* Based on visual inspection of the percent correct in healthy control performance, we graded the 12 items from easiest to hardest. We then formed three variables: ‘easy group’, containing the 4 easiest items (1, 7, 2, 4), ‘medium group’, containing the next 4 items (3, 5, 6, 10) and ‘hard group’, containing the 4 hardest items (12, 9, 8, 11). We calculated each patient’s score for the three variables (0-4). We compared the performance of the LF, RF and Controls on the three groups to investigate differences in performance based on item difficulty.
c. *Multiple-demand network* We compared overall performance in patients with versus without MD damage, controlling for age and NART (see neuroimaging investigations). We used both frequentist and Bayesian linear regression to investigate whether extent of MD involvement predicted APM performance, over and above that predicted by age and NART.

### Neuroimaging investigations

Imaging data were available for 176 patients (n=173 MRI, n=3 CT; n=110 Frontal, n=66 Non-Frontal). MRI scans were obtained on either a 3T or 1.5T Siemen scanners following a diversity of clinically-determined protocols outside our control. CT studies were obtained on Toshiba or Siemens spiral scanners. Note that since the input to the imaging models is *not* raw image data but comparatively large, manually-traced, binary lesion masks, in keeping with established practice in the field we made the assumption that the effect of variations in acquisition parameters is likely negligible and need not be explicitly modelled. Lesions were traced and independently classified using MIPAV (https://mipav.cit.nih.gov/) by JM, EC and checked by PN, who was blind to the study results. In tumour patients, the segmented lesion included the surgical cavity. The lesion masks were non-linearly normalised to Montreal Neurological Institute (MNI) stereotaxic space at 2×2×2mm resolution using SPM-12 software (http://www.fil.ion.ucl.ac.uk^79^). The lesion distribution is displayed in Figure 1. Involvement of the MD was established by comparing each patient’s normalised lesion mask with a template of MD regions in MNI space kindly provided by Professor Duncan’s group.^80^ For each patient, we determined whether their lesion involved the MD and calculated extent of MD involvement (i.e. MD lesion volume/total lesion volume x 100).

**Figure 1.**
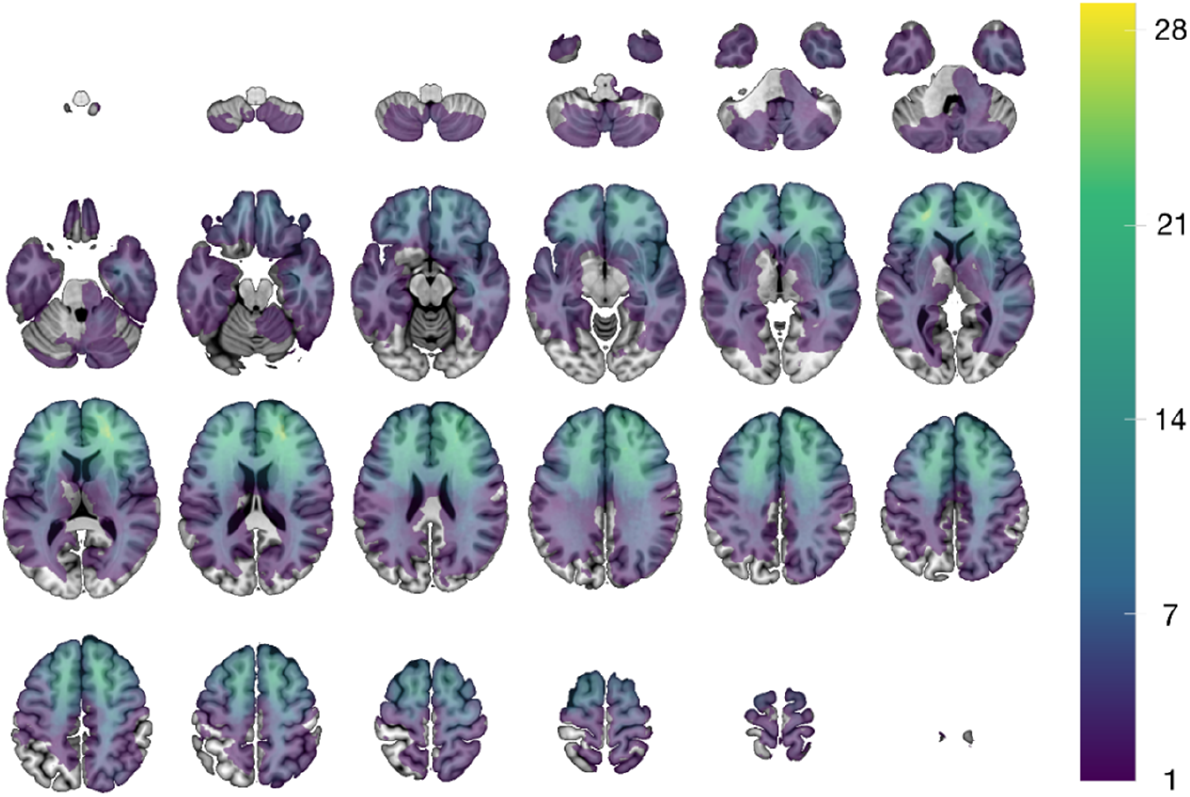
Voxel-wise sum of the 221 modelled lesions. The underlay is the SPM152 T1 template distributed with MRIcroGL(https://www.nitrc.org/projects/mricrogl). The images are displayed in neurological convention.

### Behavioural analysis

All statistical analyses were conducted using SPSS version 25. Neuropsychological data were assessed for skewness and kurtosis and tested for normality using the Shapiro-Wilk test.

One-way univariate analysis of variance (ANOVA), independent samples *t*-tests or chi-square analyses were conducted for continuous and categorical data respectively to investigate differences between Frontal, Non-Frontal and Control participants on age, gender, aetiology, chronicity, lesion volume, years of education, and neuropsychological variables (NART IQ, GNT, Incomplete Letters, S fluency and Hayling suppression errors). Following significant differences, post-hoc tests with Bonferroni correction (0.05/3=p=0.016) compared Frontal versus Non-Frontal, Frontal versus Control and Non-Frontal versus Control. LF and RF were also compared on all demographic and neuropsychological variables using t-tests.

*Standard* and *Lateralisation* analyses were performed on APM overall performance. In the *Standard* analysis we established the sensitivity and specificity of the APM to the frontal lobes. This analysis was critical because, only if there was a significant frontal deficit compared to Controls the subsequent *Lateralization* analysis was carried out to investigate unilateral left and/or right frontal contributions to APM.

In the *standard analysis*, ANCOVA was used to compare Frontal vs Non-Frontal vs Control, adjusting for age and NART. Following significant differences, post-hoc tests with Bonferroni correction (0.05/3=p=0.016) compared Frontal vs Non-Frontal, Frontal vs Control and Non-Frontal vs Control. In the *laterality analysis*, ANCOVA was used to compare LF vs RF vs LNF vs RNF vs Control, adjusting for age and NART. Following significant differences, pairwise comparisons with Bonferroni correction (0.05/4 yields p=0.0125) compared each patient group against Control (i.e. LF vs Control, RF vs Control, LNF vs Control, RNF vs Control), LF with RF and LNF with RNF.

To investigate potential differences in performance according to item difficulty in Frontal patients, we used a 3 × 3 ANCOVA with Difficulty (Easy, Medium, Hard) as the within group factor and Group (LF, RF and Controls) as the between group factor, covaried for age and NART. Significant main effects of Group were followed by simple effects analyses with Bonferroni correction.

To investigate the contribution of the MD to overall performance one-way ANCOVA was used to compare patients with (*N* = 153) versus without (*N* = 23) MD lesions, while adjusting for age and NART. We also performed a multiple linear regression analysis, using the enter method, entering APM performance as the outcome variable and age, NART and extent of MD involvement as predictor variables.

### Neuroimaging analysis

Lesion-deficit inference is complicated by the presence of correlations across damaged voxels, not just functionally—arising from a distributed neural substrate—but also pathologically—arising from the structure of the underlying pathological process^71^. Without explicit modelling of regional interactions within high-dimensional models that demand large-scale data, spatial inferences are likely to be unquantifiably distorted. In the absence of an established approach applicable to the comparatively small-scale data regimes inevitable in neuropsychology, we applied multiple inferential methods, focusing on the graph-based approach with the strongest theoretical foundations.

### Parcel-based analysis

*PLSM ANALYSES* were completed using the NiiStat toolbox for Matlab (http://www.nitrc.org/projects/niistat). To increase statistical power the brain was regionally parcellated following the JHU-MNI atlas^81^ into 189 regions of interest (ROI) spanning both grey and white matter. To assure statistical power, only ROIs with damage in >=10 patients were included. Three Freedman-Lane permutations^82^ were performed with age, NART and lesion volume always entered as nuisance regressors. Permutation thresholding (5,000 permutations) was used to correct for multiple comparisons and control the family-wise error rate. An alpha of 0.05 was the threshold for significance. To investigate the contribution of the MD to APM overall performance, PLSM analyses were repeated with age, NART and proportion of MD involvement entered as nuisance regressors. PLSM analyses were conducted on Hayling suppression errors (scaled scores), with age, NART and lesion volume or age, NART and proportion of MD involvement entered as nuisance regressors.

### Bayesian multivariate lesion-deficit modelling of MD dependence

To quantify the regional contribution of components of the MD, Bayesian multivariate regression implemented in BayesReg v1.91 was performed with each connected component of the MD map treated as a predictor variable, and age and NART added as nuisance covariates. A selection of shrinkage priors (ridge, lasso, g, horseshoe, horseshoe+) and noise models (Gaussian, Laplace, Student t distribution) were evaluated, choosing g and Student t based on the widely applicable information criterion (WAIC): a standard interpretable metric for Bayesian model comparison.^83^ The posterior distributions of the regression coefficients were estimated with Markov chain Monte Carlo sampling over 100000 samples with a 100000 burn-in interval and thinning set at 10, reporting the means and standard deviations of the regression coefficients that survive a 95% Bayesian credibility interval. The effective sample size was >97 for all models.

### Graph lesion-deficit modelling

Where the neural support of a function is distributed across a set of connected regions, the optimal way of identifying it is through explicit modelling of both anatomical locations and their interactions. Even where the neural support is local, the structure of the lesion pathology used to reveal it need not be, and itself requires modelling of distributed relations. By structure here is meant characteristic spatial patterns of coincident damage dictated by the underlying pathological process, such as the patterns of ischaemic damage the vascular tree enforces in stroke. The difficulty is amplified when both the neural and the pathological are distributed, for the former must then be disentangled from the latter: a problem for which there is no established solution. Here we adopt an approach based on statistical models of graphs. The fundamental idea is to conceive the brain as a densely interconnected graph, where each node is an anatomical location and each edge indexes the extent to which its connected nodes share a set of properties. In the context of lesion-deficit mapping, the properties of interest are the presence of damage, the associated deficit, and nuisance factors that could confound their relations. First, we apply conventional network-based statistics, fitting a general linear model to APM scores, revealing a network of dependence driven jointly by functional anatomy and spatial patterns of damage. Second, we exploit recent developments in Bayesian stochastic block modelling to identify communities of voxels distinctively influenced by fluid intelligence, disentangled from the incidental spatial structure of lesions.

### Network based statistics

The non-linearly registered lesion masks were linearly resampled to a resolution of 12 mm^3^. This resolution offers considerably finer anatomical detail than published parcellation schemes^84–87^ and avoids the potentially biasing effects of their structuring determinants. We provide a bar plot showing the mean node volume (± 95% confidence interval) if our approaches compared with commonly used parcellation schemes in Supplementary Figure 1.

A graph where voxels are the *nodes* and their adjacent neighbours the *edges* was created as an adjacency matrix, labelling any edge that linked two lesioned nodes as 1 and all others as 0. This process yielded a graph of order and size 1017 nodes and 516636 edges for each patient in which the lesion was larger than a single 12mm^3^ voxel (N=172). The choice of a 12mm^3^ voxel size was constrained by the tractability of the statistical model, in line with the practice of others in related domains.^23, 88^

We proceeded to model lesion adjacency matrices with the Network-Based Statistics connectome toolbox (v1.2).^90^ NBS is an established statistical framework for network analysis, described in extensive detail elsewhere^91–92^. In brief, it implements a non-parametric approach to mass-univariate statistical inference on the edges of large graphs, yielding family-wise error (FWER)-corrected *p*-values for each edge via permutation testing^93^. The approach can be viewed as the graph analogue of the mass-univariate voxel-wise methods familiar from functional imaging and voxel-based morphometry. It has been widely applied to investigate the organisation of brain networks.^90–92, 94^

Here the inputs were the lesion graphs of each patient, with APM as the predictor and NART and age as nuisance covariates. The model was fitted with 50,000 permutations, with a criterion for statistical significance set at family-wise error rate corrected *p*<0.05, yielding an inferred group-level network significantly associated with Gf. We evaluated the community structure of this inferred network—the presence of clusters of voxels defined by similar inferred connectivity—with a Bayesian, weighted, non-parametric, hierarchical, generative stochastic block model,^95, 96^ with additional simulated annealing to approximate the global optimum of the function (see below). Edges were weighted by the significant t-statistic adjacency matrix from the NBS model. To examine the potential influence of aetiology, we compared this NBS model to another identically configured except for the addition of aetiology as a nuisance covariate (see Supplementary material).

To illustrate the relation between the inferred network and fluid intelligence, we created a set of Bayesian regression models with a target of APM adjusted for NART, and predictors constructed— across separate models—from the dichotomized overlap between a lesion and the NBS-identified network, or from the number of nodes of each patient’s graph included in the NBS network. We also ran a multivariate regression model with the lesion adjacency matrix as columns of predictors. We evaluated all models with various prior shrinkage schemes, using the WAIC to select the most appropriate prior distributions and model goodness-of-fit.^83, 97–98^ All regression models were implemented in BayesReg v1.2, and employed a burn-in of 50000, taking 100000 samples from the posterior distribution within a single MCMC chain. Note these analyses are not independent and are designed to be merely illustrative of the NBS model from which they are derived.

### Bayesian hierarchical stochastic block modelling

The foregoing simple network-based statistical model is potentially confounded by the anatomical structure of pathological damage. It is also tractable only at relatively coarse spatial resolutions. To overcome these defects, we exploited a recent innovation in the statistical modelling of graphs: Bayesian stochastic block models.^103^ These are nonparametric probabilistic statistical models of the network structure of graphs that enable robust inference to distinct patterns of connectivity arising as network “blocks” or “communities” within them. In the context of a graph model of the lesioned brain, such communities may be shaped by the neural substrate of the behaviour under study, the anatomical patterns of damage, or an interaction between the two. The approach allows us to disentangle these two distinct types of node connectivity, in our case isolating the neural dependents of APM performance from the incidental structure of the lesions used to reveal them. We employ a specific kind of Bayesian stochastic block model designed to incorporate layered, multiple attribute properties.^103^ The layered formulation enables robust inference to the separability of the two types of connectivity by Bayesian model comparison of variants whose layered structure either respects or ignores them^95^. Though a comparatively recent innovation, such models rely on well-established principles of Bayesian inference and graph theory, and are underwritten by their theoretically proven validity^95, 100–102^.

Graph theory provides a powerful method of modelling complex systems that combines flexibility with intelligibility.^92^ It treats individual factors of interest as the “nodes” of a network, and their interactions as the connections, or “edges”, between them. In the context of lesion deficit inference, the nodes identify anatomical locations in brain, and the edges describe their pairwise relations. Two nodes may be related by their association with a deficit when lesioned, or by their tendency to be involved in the same lesion, regardless of the deficit. The former is the effect of interest, the latter is a potential confounder we wish to eliminate. To disentangle the two forms of relation we create a layered, weighted, undirected graph whose layers correspond to the two different kinds of association. Confining each form of relation to its own layer compels the model to disentangle them in inferring the community structure of the graph. We can compare a layered model of this kind to a null model where the edges are randomized across layers, employing Bayesian model comparison based on the minimum description length of the model. Finding the layered model superior to the null is evidence of the successful separation of the structuring effects of APM and lesion co-occurrence we seek here. A detailed exposition of the inferential approach is given in Supplementary material.

To model our data, each non-linearly registered lesion was resampled to 4mm^3^ resolution, and the lesion adjacency matrix constructed for each patient as before. This resolution is much finer for than conventional parcellation schemes published in the wider literature schemes (Supplementary Figure 1).^84–87^ We then constructed an undirected, weighted graph combining all individual lesion networks across all patients. This network comprised nodes corresponding to all voxels of the brain, and edges between voxels adjacent. These edges were weighted by two variables: the count of the number of times a voxel and an adjacent neighbour were damaged *together* —a *lesion co-occurrence weight*— and the inverse of the patient’s APM score divided by NART—an adjusted *APM weight.* Naturally, the graph was undirected, as the direction of any relationship between collaterally lesioned areas is not informed by the data at hand.

We filtered edges to limit analysis to the top 50% connected nodes, removing edges with fewer than ∼3 connections, where sampling was too low to support robust inference, but permitting still full brain coverage. This yielded a graph of order and size 27509 nodes and 285545 edges. There were no node self-loops. We rescaled both lesion co-occurrence and APM edge weights to the range 0 to 1.

We proceeded to evaluate the community structure of this network with a non-parametric Bayesian hierarchical weighted stochastic block model incorporating layered and attributed properties^103, 109^ implemented in graph-tool (https://graph-tool.skewed.de).^95–96, 102^ We began by fitting a null model, with the two kinds of edge weight—adjusted APM and lesion co-occurrence—randomly distributed across two layers. We then fitted a test model with each type of weight consistently assigned to its own layer. Adjusted APM weights were modelled as Gaussian; lesion co-occurrence weights as Poisson distributions. Having initialised a fit, we used simulated annealing to further optimise it, with a default inverse temperature of 1 to 10.^96^ We did not specify a finite number of draws, rather we specified a wait step of 100 iterations for a record-breaking event, to ensure that equilibration was driven by changes in the entropy criterion, instead of driven by a finite number of iterations^102^

We used model entropy to determine if the layered model fit was better than the null, indicating that the inferred community structure distinguished APM and lesion co-occurrence effects. To visualise the inferred communities, we backprojected the incident edge weights onto the brain, deriving the mean and 95% credible intervals for comparison. To examine if modelling lesion co-occurrence requires explicit consideration of aetiology, we replicated the model with the addition of aetiology as a third layer, again conducting formal comparison against a randomised null (see Supplementary material).

### Synthetic ground truth evaluation

The substrate of a function is definitionally unknown: it is what we are seeking to infer. To examine the comparative fidelity of a set of models we therefore need synthetic ground truths^72^ of the complexity likely to obtain in reality. Here we used the meta-analytic repository NeuroQuery^104^ to create six realistically complex and distributed ground truth maps across the domains of action, aversion, language, mood, motor and sensation (see Supplementary materials). The intersection between each lesion and each ground truth was then used to generate a hypothetical deficit for each patient and each domain, and the stochastic block model was subsequently applied exactly as in the case of the real data. The fidelity of the inferred maps was then quantified by their Dice score, and compared to a standard mass-univariate voxel-based lesion-deficit mapping baseline (see Supplementary materials).

### Data and code availability

The data and code that support the findings of this study are available from the corresponding author, LC, upon reasonable request.

## Results

### Demographic and behavioural investigations

Frontal, Non-Frontal and Control patients were well-matched for age, gender, chronicity and years of education (all *p* > 0.05). There was no significant difference in lesion volume between LF and RF or between LNF and RNF. There was a significantly greater proportion of tumour patients in Frontal than Non-Frontal (χ2 (1, N = 227) = 5.68, p < 0.05) and a significantly greater proportion of stroke patients in Non-Frontal than Frontal (χ2 (1, N = 227) = 7.05, p < 0.01). However, there were no significant differences in the proportion of tumour or stroke patients between LF and RF (all *p* > 0.05) or between LNF and RNF (all *p* > 0.05).

There were no significant differences between Frontal, Non-Frontal and Control participants for NART, IL or GNT scores (all *p* > 0.05; see Table 1). One-way ANOVAs found highly significant differences between Frontal, Non-Frontal and Control participants for S fluency and Hayling suppression errors (*F*(2,238) = 25.319; *p* < .001; *F*(2,192) = 21.266; *p* < .001, respectively). Post-hoc tests showed Frontal performed significantly worse than Non-Frontal and Control on S fluency (*p<*.001; *p* <.001, respectively) and Hayling suppression errors (*p*<.001; *p* <.001, respectively). Pairwise comparisons revealed that LF were significantly more impaired than RF on S fluency (*p*<.01), RF were significantly more impaired than LF on Hayling suppression errors (*p*<.05; Table 1).

### Overall performance

*Standard analysis*: A one-way ANCOVA controlling for age and NART, found a highly significant difference between Frontal, Non-Frontal and Control participants in overall performance (*F*(2,387) = 18.491; *p* < .001). Post-hoc tests showed that Frontal performed significantly worse than Non-Frontal (*p* < .01) and Control (*p* <.001). There was no significant difference between Non-Frontal and Control (corrected p =.185; Table 2)

**Table 2.** Overall performance on APM. n = Number. SD = Standard Deviation. Scores with significant p values are in bold. **= p <0.01; ***= p <0.001. ^a^indicates significant difference from non-frontal patients. ^b^ indicates significant difference from healthy control participants. ^c^ indicates significant difference between left and right frontal patients.

| Performance | Healthy control | Frontal | Non-Frontal | Frontal |  | Non-Frontal |  |
| --- | --- | --- | --- | --- | --- | --- | --- |
| | | | | Left<br>$n = 69$ | Right<br>$n = 77$ | Left<br>$n = 39$ | Right<br>$n = 42$ |
| Mean number correct /12 (SD) | 8.67<br>(2.41) | **7.07<sup>a\*\*</sup> b\*\*\***<br>(2.78) | 8.07<br>(2.09) | **7.71<sup>b\*\*</sup>**<br>(2.49) | **6.49<sup>b\*\*\*</sup> c\*\***<br>(2.93) | 8.00<br>(1.92) | 8.14<br>(2.26) |

*Lateralization analysis:* A one way ANCOVA controlling for age and NART found a highly significant difference between LF, RF, LNF, RNF and Control participants (*F*(4,385) = 12.237; *p* < .001). Pairwise comparisons showed a significant difference between RF and Controls (*p*<.001) and LF and Control (*p*<.01). Importantly, RF were significantly more impaired than LF (*p*<.01). There was no significant difference between RNF and Control or LNF and Control; table 2). Notably, performance fell <1.5 standard deviations below Control in 43% of RF but only in 22% of LF.

### Item difficulty

A 3×3 ANCOVA, controlling for age and NART, revealed a significant main effect of difficulty (F(2,364) = 5.360; p=0.005) and Group (F(2,182) = 22.707; p < .001). Critically, there was also a significant interaction between difficulty and group (F(2,364) = 10.822; p<0.001). Post-hoc pairwise comparisons showed significant differences for the medium group between RF and Control (p<.001), LF and Control (p<.05), and RF and LF (p<0.05). Thus, Frontal patients were worse than Control, and RF performed the poorest. For the hardest group there were significant differences only between RF and Control (p<.001), and LF and Control (p<.001). There were no significant differences for the easy group. Thus, RF impairment on the APM appears to be driven by poor performance on the medium items. Closer inspection revealed that three specific items (3, 5 and 6) were responsible for driving the RF poorer performance than LF, with an accuracy decrement of more than 20% in RF.

*Multiple-demand network* A one-way ANCOVA, controlling for age and NART, showed no significant difference in overall performance between patients with vs without MD lesions (*F*(1, 172) = 1.88, *p =* .172). A linear regression analysis, with age, NART and extent of MD involvement entered as predictor variables, significantly predicted APM performance (r2 = .28, *F*(3, 172) = 21.726, p < .001). However, only age and NART (both p<.001) were significant predictors. Extent of MD involvement did not significantly contribute (p = .410).

### Parcel-based analysis and Bayesian multivariate analysis of MD

PLSM analyses with age, NART and lesion volume entered as nuisance regressors, revealed that poorer overall performance was associated with right posterior middle frontal gyrus, pars opercularis, precentral gyrus, superior corona radiata and external capsule lesions. When the proportion of MD involvement was entered as a nuisance regressor instead of lesion volume the results remained unchanged.

PLSM analyses on Hayling suppression errors, with age, NART and lesion volume entered as nuisance regressors revealed that poorer performance was associated with right posterior middle frontal gyrus and pars opercularis lesions. When the proportion of MD involvement was entered as a nuisance regressor instead of lesion volume the results remained unchanged.

Bayesian multivariate modelling of individual MD components yielded as credibly predictive only NART (posterior mean coefficient 0.484, 95% credibility interval 0.336 to 0.630), age (mean -0.321, 95%CI -0.455–-0.187), and a right-sided MD component falling within precentral and posterior medial frontal gyrus (mean -0.310, 95% CI -0.587 to -0.021). The credibility intervals of the coefficients of other MD components all crossed zero.

### Network lesion-deficit modelling

Network-based statistics identified a distinct predominantly right frontal network associated with reduced APM (FWER-corrected *p*<0.0001, t-thresh >3.1) (Figure 2). The regions with the greatest number of significant nodes (in order of descending degree count) included the right superior frontal gyrus (degree count 22), right middle frontal gyrus (17), right frontal pole (16), right anterior cingulate cortex (16), left superior frontal gyrus (6), right inferior frontal gyrus (4), left anterior cingulate cortex (4), right caudate nucleus (3), right mid cingulate cortex (3), right precentral gyrus (2), right juxtapositional lobule (1), right frontal operculum (1) and right anterior insula (1). A stochastic block model partitioned the network into a structure with 3 clustered components, broadly encompassing medial wall, superolateral cortical surface and a superior frontal gyrus-dominant component. The addition of lesion aetiology to the list of covariates in the NBS model yielded a near-identical result (test-statistic correlation between significant edges, r .99, *p*<0.0001) (Supplementary Figure 2).

**Figure 2.**
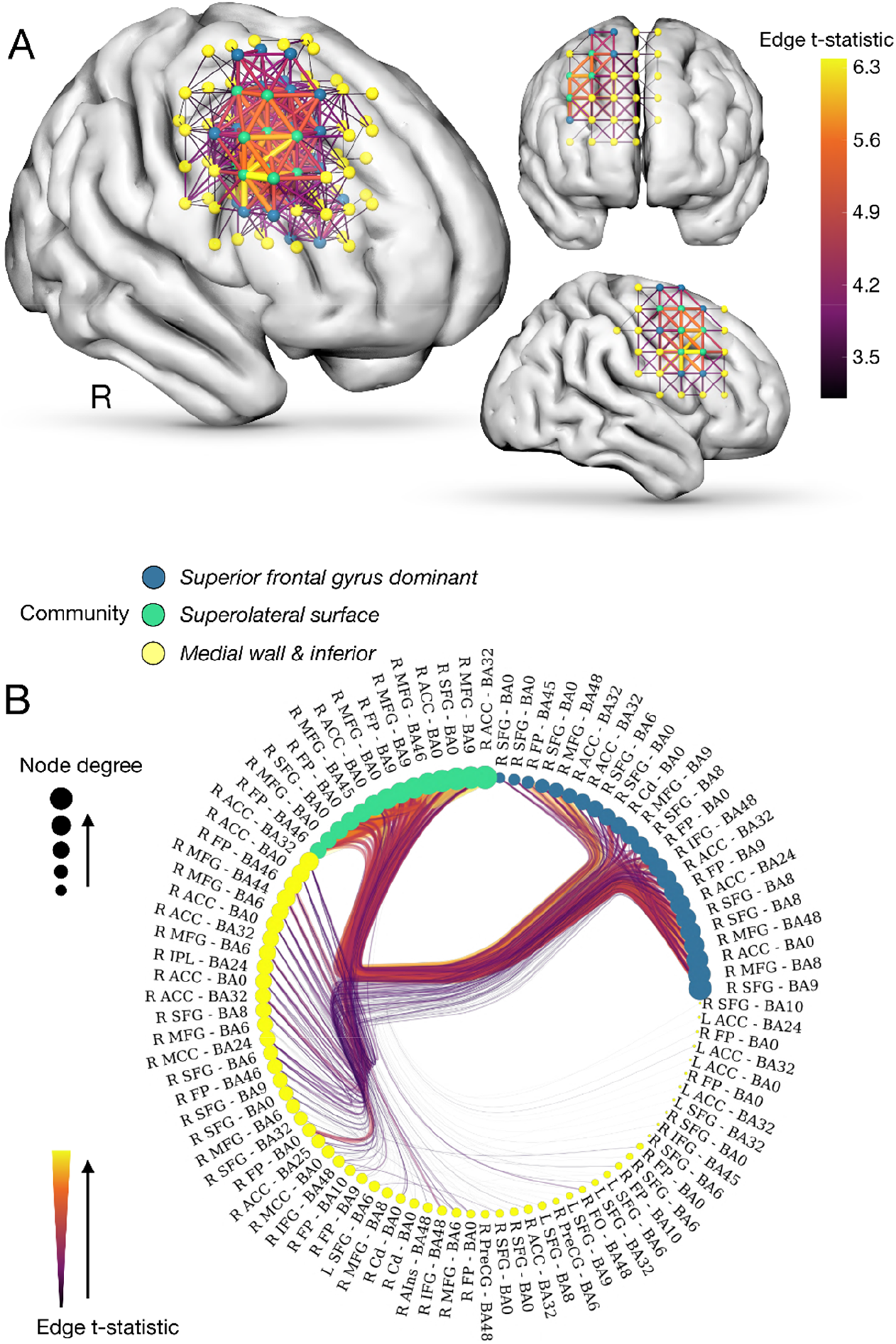
Network-based statistic maps. **A.** Network-based statistics identify a significant network associated reduced adjusted APM scores (FWER-*p*<0.0001). **B.** Radial graph of the community structure of the network inferred from a stochastic block model of its statistics shows that the network clusters into three discrete components encompassing the superolateral cortical surface, the medial (and inferior) wall and a superior frontal gyrus dominant cluster. Nodes are colour-coded in accordance with their stochastic block model cluster. Node size is proportional to node degree count. Edge width and colour is proportional to the t-statistic from the model, with a thicker and more yellow line denoting a stronger link between a given network connection and a reduced adjusted APM score. Abbreviations: ACG, anterior cingulate gyrus; L, left; IFG, inferior frontal gyrus; IFG-pt, inferior frontal gyrus pars triangularis; MFG, middle frontal gyrus; OFC, orbitofrontal cortex; PreCg, pre-central gyrus; PoCg, post-central gyrus; R, right; SFG, superior frontal gyrus; SPL, superior parietal lobule.

Bayesian univariate regression analysis confirmed that patients with lesions overlapping with the network exhibited significantly lower adjusted APM scores, (R^2^ 0.105, coefficient mean ± SD -0.265 ± 0.068 (95% credible interval [CI] -0.396 to -0.129) (Figure 3). The extent of overlap, indexed by the degree (i.e. number of nodes) shared between an individual lesion network and the inferred network exhibited a strong log-linear relationship to adjusted APM (R^2^ 0.190, coefficient mean ± SD -0.002 ± 0.0005 (95% CI -0.00250 to -0.00041). Bayesian multivariate regression models of adjusted APM predicted by the lesion network adjacency matrix yielded a fit with R^2^ 0.640. It is important to note that these regression analyses are not independent of the NBS model: they do not provide further evidence but rather qualify its fidelity.

**Figure 3.**
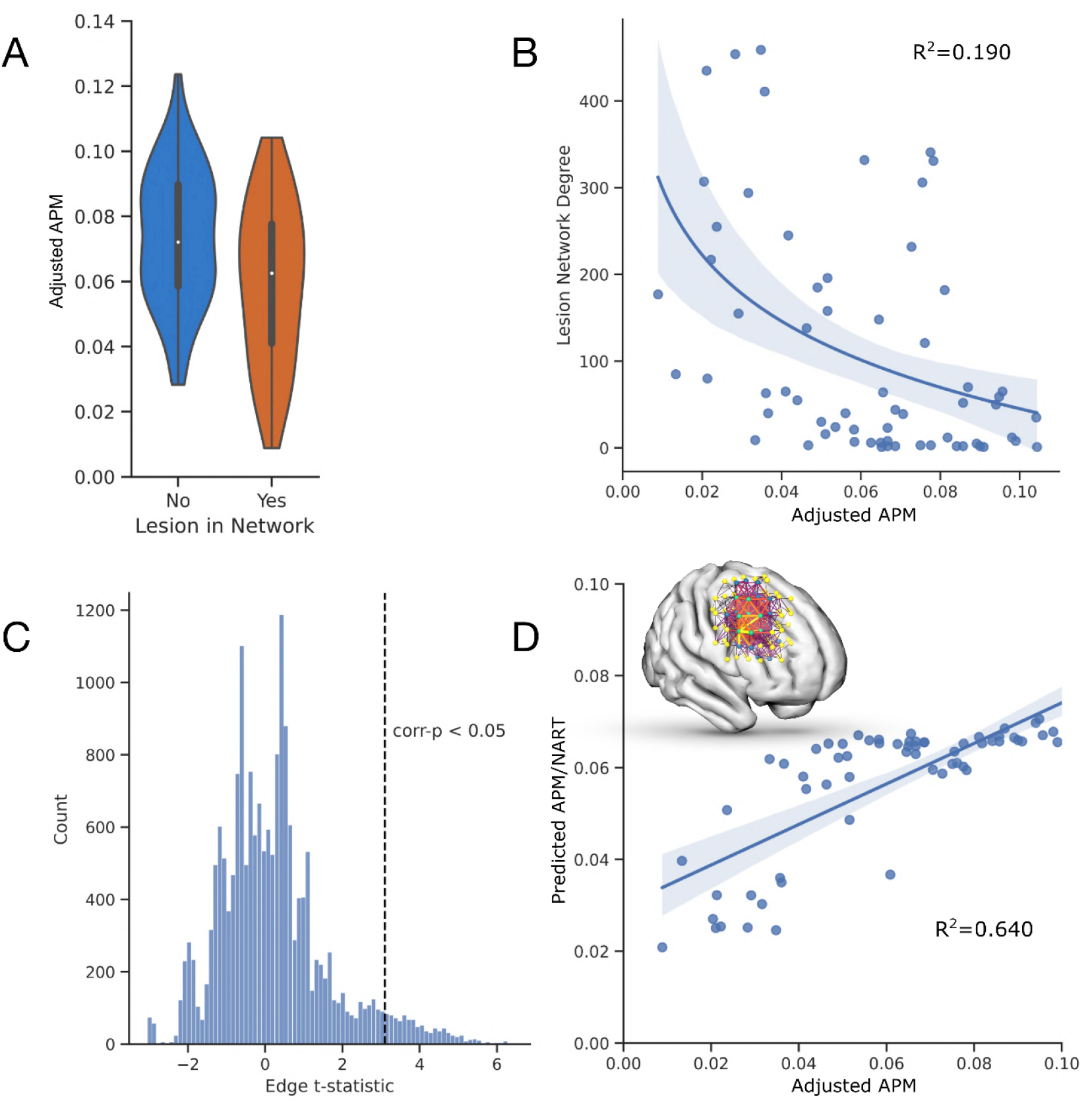
Network-based statistic behavioural correlations. **A.** Violin plots of the adjusted APM scores of patients whose lesions do or do not overlap with the inferred network illustrate significantly lower APM scores in the former (R2 0.105, coefficient mean ± SD -0.265 ± 0.068 (95% credible interval [CI] -0.396 to -0.129). **B.** Scatter and line plot shows that the degree count of the overlap of a lesion with the inferred network significantly correlates with adjusted APM scores within a univariate Bayesian regression model (R^2^ 0.190, coefficient mean ± SD -0.002 ± 0.0005 (95% CI -0.00250 to -0.00041). **C.** Histogram of the edge t-statistics from the network model illustrates the population of edges significantly associated with the APM after multiple comparisons correction. **D.** Scatter and line plot shows the predictability of adjusted APM from the network adjacency matrix within a multivariate Bayesian model (R^2^ 0.640).

### Generative hierarchical stochastic block modelling of APM performance

The foregoing models inevitably conflate the distributed spatial structure of the underlying neural dependence with that of the causal pathology. To disentangle the two, we need a network model capable of separating the target effects of APM performance from the incidental effects of lesion co-occurrence. This can be achieved with a layered nested stochastic block model, where adjusted APM and lesion co-occurrence weights are distributed in two distinct layers, yielding layer-specific patterns of community structure reflecting the distinct effect of each weight on the network. This model achieved substantially lower entropy—881118.22 vs 1182697.66 nats—than a null model with weights randomised across the two layers (Figure 4), providing inferential support for distinguishing adjusted APM from co-occurrence effects. This translates to a posterior Odds ratio of the layered formulation being e^301579^ more likely than the non-layered null.

**Figure 4.**
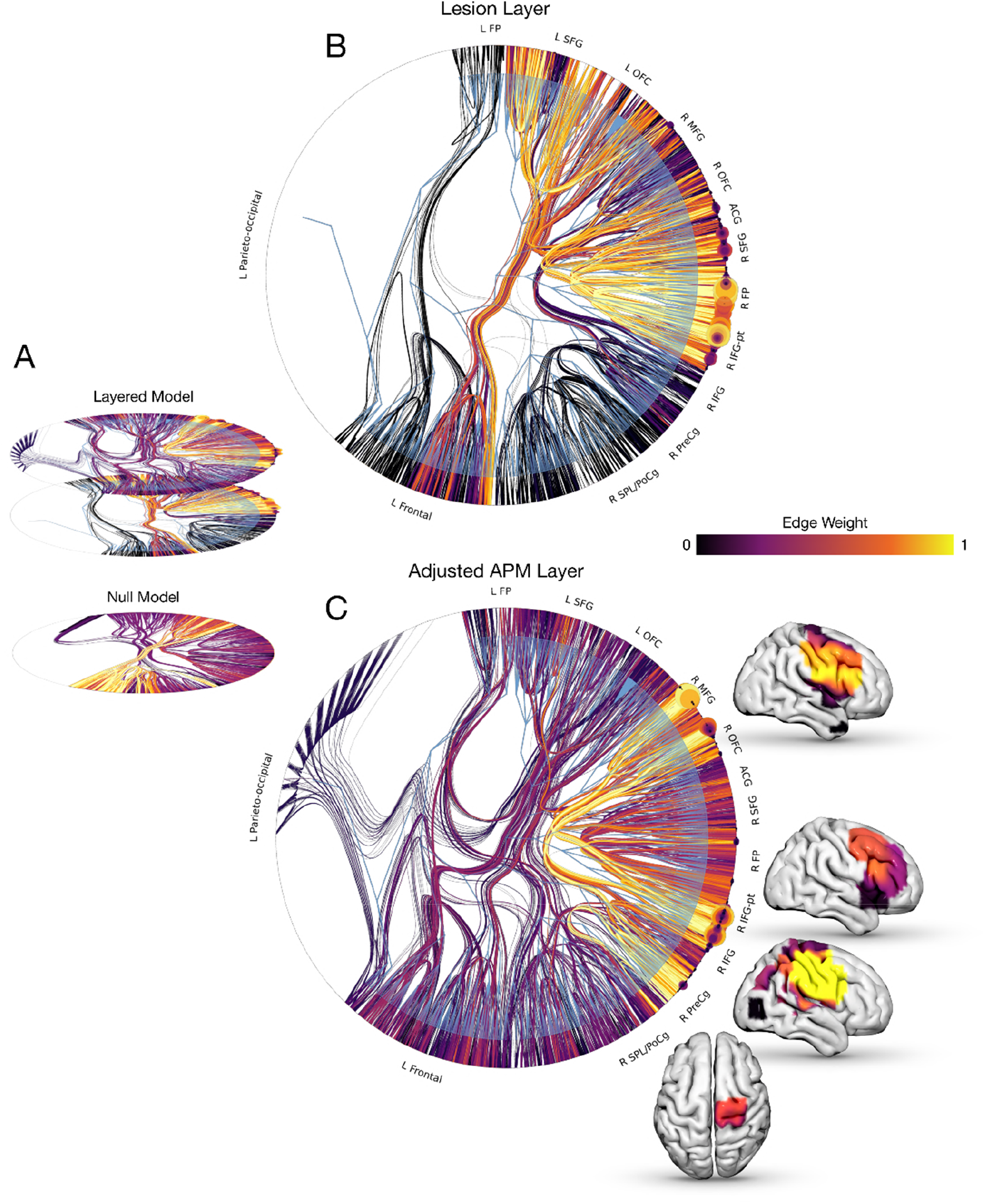
Stochastic block model lesion-deficit mapping. **A.** Radial graphs of stochastic block models with adjusted APM and lesion co-occurrence layered (top), vs randomly distributed across layers (bottom). Edge colour and width is proportional to the associated edge weight. Model entropy favoured the layered over the null model. **B.** Radial graph illustrating the layered stochastic block model fit with edge colour and width proportional to the lesion co-occurrence weight, and node colour and size proportional to the lesion-weight degree. This demonstrates a community of highly interconnected voxels involving the bilateral frontal pole and orbitofrontal cortex, right superior and inferior frontal gyrus and anterior cingulate gyrus. **C.** Radial graph illustrating the layered stochastic block model fit with edge colour and width proportional to the adjusted APM weight, node colour and size proportional to the APM-weight degree. This illustrates a characteristically different segregation of brain communities, with high edge incidence linking the right middle and inferior frontal gyrus, (including pars triangularis), right pre-central gyrus and right superior parietal lobule. Brain images are overlayed corresponding to the posterior mean edge weight at these communities.

**Figure 5.**
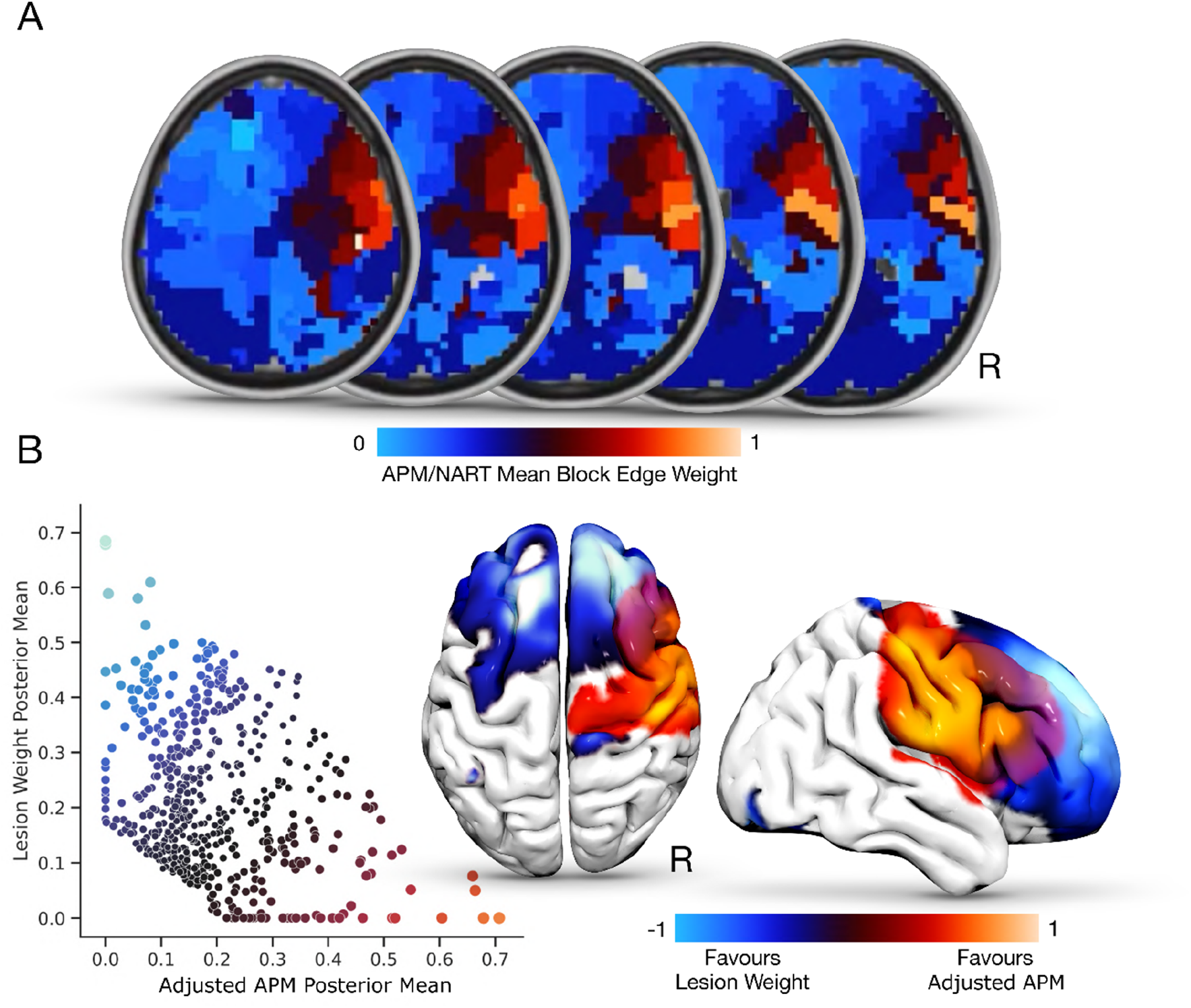
Graph community properties. **A.** Axial slices of the mean posterior edge weight for each block at the l1 aggregation, with more red-orange areas corresponding to a greater value and greater relation to adjusted APM. **B.** Scatterplot illustrating the relationship between posterior mean edge weights at each community block, for both the lesion weight (y-axis) and adjusted APM (x-axis), with brain reconstructions overlaying these findings. Of note, bilateral frontal-based blocks depicted higher lesion-weight edges, with right fronto blocks more implicating APM. Abbreviations: ACG, anterior cingulate gyrus; L, left; IFG, inferior frontal gyrus; IFG-pt, inferior frontal gyrus pars triangularis; MFG, middle frontal gyrus; OFC, orbitofrontal cortex; PreCg, pre-central gyrus; PoCg, post-central gyrus; R, right; SFG, superior frontal gyrus; SPL, superior parietal lobule.

The community structure was composed of blocks dominated by adjusted APM, lesion co-occurrence, or neither weight. The adjusted APM layer revealed a set of brain communities with high edge incidence linking the right middle and inferior frontal gyrus (including pars triangularis), right pre- and post-central gyri, and—weakly—the right superior parietal lobule. These communities were sharply distinct from the lesion weight.

Many of the spatial constraints on the configuration of lesion patterns are imposed by the basic anatomy of the brain and will be shared across aetiologies; those that are not will arise as additional heterogeneity the stochastic block model could theoretically absorb. To determine if aetiology has a substantial structuring effect that merits explicit accounting, we reran the model with an additional, third layer identifying the aetiology of each lesion. This model exhibited far greater description length (2133947.48), 2.4x that of the above for an increase of this single feature, indicating a poorer fit to the data and providing no grounds for preference over the simpler model. The anatomical pattern of APM-sensitive communities was in any event very similar (Supplementary Figure 3).

### Synthetic ground truth evaluation

Bayesian model comparison showed all layered models to be more plausible than the null (Supplementary Figure 4). Compared with VLSM, the stochastic block model achieved significantly superior results across all domains (*p*=0.028) (see Supplementary materials) (Supplementary Figure 5). These experiments also demonstrated qualitatively the tendency of VLSM to mislocalise in response to the underlying lesion structure, and the ability of the stochastic block model to resist it.

## Discussion

Our study represents the first large-scale investigation of the distributed neural substrates of fluid intelligence in the focally injured brain. We investigated one of the most widely used Gf tests, the APM, in the largest number of patients with single, focal, unilateral, right or left, frontal or non-frontal lesions and controls. We analysed overall performance, item difficulty and the contribution of MD involvement. For the first time, non-parametric Bayesian stochastic block models were used to reveal the intricate community graph structure of lesion deficit networks, disentangling functional from confounding pathological distributed effects.

Similar to other groups^21, 105–107^ and in-keeping with our previous studies^71^ we adopted a mixed aetiology approach. Previous comparison of a large frontal and non-frontal sample with different aetiologies on the APM and other executive tests showed that aetiology was not a strong predictor of frontal or non-frontal deficits.^9, 108^ Hence, different aetiologies do not result in more severe impairments than others and combining across vascular and tumour pathologies is unlikely to significantly distort neuropsychological performance.^76^ Instead, focal lesions may relate more closely to the region of damage rather than aetiology. Moreover, data from multiple aetiologies will tend to attenuate distorting effects arising from pathologically driven characteristic patterns of lesion co-occurrence that are widely recognized to bedevil both network and focal lesion-deficit studies. Indeed, less spatial distortion caused by the structure of the pathology may be expected if multiple pathologies differing in their spatial properties are used.

Though Gf is widely thought to be dependent on the integrity of the frontal lobes, only a handful of focal lesion studies, based on modest samples, have found impairments in following frontal lesions.^9, 21, 55^ Applying an array of lesion-deficit models to large scale data, we found APM performance to be specifically vulnerable to the integrity of the right frontal lobe, and largely resistant to damage elsewhere. The left frontal lobe appears to make a contribution to APM performance, if a more modest one. We found that the performance of the left frontal patients was significantly different from healthy controls and non-frontal patients. However, the left frontal patients performed significantly better than the right frontal patients did.

Our findings speak to the theories of non-frontal involvement in Gf. The proponents of P-FIT have argued that impairment of Gf should follow lesions of the posterior and anterior regions that putatively subserve it.^23, 24^ We found no evidence of such non-frontal causal dependence on APM performance. It is possible that functional imaging findings merely reflect a correlation between Gf and posterior areas non-critically engaged by the necessary perceptual input.

Our results are also relevant for the MD proposal. Notably, Woolgar *et al*.^80^ investigated 80 patients with cortical lesions with a Gf task (Cattell Culture Fair IQ test). Though the authors reported a significant correlation between MD involvement and Gf performance overall, in the 44 patients with purely frontal damage the relationship was not significant when non-MD lesion volume was taken into account. So, as far as the frontal lobes are concerned, the authors’ theoretical claim was not strongly empirically supported. In our study patients with or without MD damage did not differ significantly in performance on the APM. Moreover, the extent of MD involvement did not contribute to performance. These findings do not support the claim that MD is the seat of Gf, but neither do they exclude it: absence of evidence is not evidence of absence. In this context, we note that the Woolgar et al^80^ study included 30 non-frontal patients. Of these only 7 were patients with unilateral parietal lesions, whilst 2 patients had biparietal lesion. In contrast, our non-frontal sample includes 81 patients, of which 20 have a unilateral parietal lesion. Hence, our study has far greater coverage of non-frontal MD areas than Woolgar’s study. While we cannot rule out the possibility that lower power may be a factor relevant for our conclusion regarding weak contributions from non-frontal MD lesions, our sample is larger than that of the study, which produced the opposite conclusions. Moreover, the Woolgar et al. study reported for their parietal patients that MD lesion volume was a significant predictor for performance on fluid intelligence with and without non-MD lesion volume partialled out (r=-0.65, p=0.042; r=-0.63, p=0.035 respectively). We were not able to replicate this effect in our larger group (r=-0.063, p=0.811; r=-0.17; p=0.950).

Our findings of greater involvement of the right frontal lobe in APM performance were complemented and extended by our neuroimaging analyses. Both conventional network statistics and non-parametric Bayesian stochastic block modelling heavily implicated the right frontal lobe. Crucially, this localisation was confirmed on explicitly disentangling—uniquely in the field of lesion-deficit mapping—functional from pathology-driven effects within a layered stochastic block model, prominently highlighting a right frontal network including the middle and the inferior frontal gyrus, including pars triangularis, and pre- and post-central gyri, with a comparatively weak contribution from superior parietal lobule. The marked structuring effects of lesion co-occurrence observed highlight the importance of explicitly modelling them in lesion-deficit inference, whether in the context of network or focal analysis.

Standard PLSM analyses, potentially confounded by lesion co-occurrence effects, suggested that poorer performance was associated with damage to a right frontal network including posterior middle frontal gyrus, pars opercularis, precentral gyrus, superior corona radiata and external capsule, invariantly to the degree of MD involvement. That a similar set of RF regions were implicated in Hayling suppression errors, a verbal test, suggests that function lateralization in the frontal lobes is not explained by task sensitivity to language alone.

Behaviourally we found a highly significant interaction between item difficulty and frontal lesion lateralisation. The asymmetry in performance was nearly three times greater for the middle than for the highest level of difficulty, with neither population nearing the ceiling or floor. Why might that be?

While complete agreement is lacking, factor analyses of progressive matrices indicate at least two material components. Dillon *et al*.^110^ identify a factor related to “perceiving the progression of a pattern” (p.1301), and another to “the addition and/or subtraction of elements”. Lynn *et al*.^111^ offer respectively “Gestalt continuation”, following Van der Ven and Ellis^112^, and “verbal-analytic reasoning”. It is apparent that the three medium items (3, 5, 6) of the APM showing the greatest lateralisation, are all those where perceiving the progression of a pattern is an obvious approach. By contrast, the non-ceiling items with the smallest lateralisation (10, 11, 12) are all those where addition or subtraction come into play. Though the limited number of items precludes firm conclusions on factorisation, these findings suggest the components of APM may lateralise in different ways. Inferences involving the perception of a progressive pattern may be especially sensitive to the integrity of the right frontal network.

In conclusion, our study represents the most robust investigation of the hitherto poorly characterized Gf in patients with single, focal, unilateral lesions. Our approach of combining novel graph-based lesion-deficit mapping with detailed investigation of APM performance in a large sample of patients provides crucial information about the neural basis of fluid intelligence. We suggest that a right frontal network, rather than a wide set of regions distributed across the brain, is critical to the high-level inferences, based on perceiving pattern progression, involved in Gf. Our findings further corroborate the clinical utility of APM in evaluating Gf and identifying right frontal lobe dysfunction.

## Supplementary Material

### Stochastic block models

A stochastic block model (SBM)^1^ is a generative model of the community structure of a graph composed of *N* nodes, divided into *B* blocks with edges *e_rs_* between blocks *r* and *s*. The model can be framed hierarchically, where edge counts *e_rs_* form block multigraphs with nodes corresponding to individual blocks and edge counts arising as edge multiplicities between block pairs, including self-loops. We seek to infer the most plausible partition {*b_i_*} of the nodes, where {*b_i_*} ∈ [*1, B*]*^N^* identifies the block membership of node in observed network *G*, with maximisation of the posterior likelihood *P*(*G*|{*b_i_*}). The result is a hierarchically organised community structure of nodes assigned into blocks that yields the most compact representation of the graph, as indexed by its minimum description length^2^, Σ. The general approach is described in further detail elsewhere^1^.

An extension to the hierarchical SBM is its layered formulation, permitting modelling of a graph structure where edges reflect disparate forms of interaction^3^. In the context of lesion deficit mapping, we can use a layered SBM to model relations between voxels driven by two distinct effects: the underlying neural dependence of the function of interest and the pathological structure of the lesions used to map it. Key here is formal comparison between models that encode these effects separately, within their own layers, vs those where the distinction is not respected. In a Bayesian setting^3^, the procedure for model selection amounts to finding the model maximising posterior likelihood as

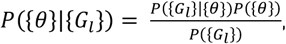

where {*θ*} denotes the shorthand for the model parameters. In our case, 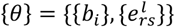, where *N* nodes are divided into *B* blocks via the membership vector {*b_i_*} ∈ [*1, B*]*^N^*, and the distribution of covariates in edges in groups *r* and *s* is given by the edge counts *e_rs_*, with 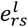 corresponding to the former at a given layer. *P*({*θ*}) is the prior probability on these parameters, with *P*({*G_l_*}) corresponding to the normalisation constant. The approach is further detailed by Peixoto^3^, formulating the most succinct representation of the data as one with the minimum description length^2^, Σ. Since the prior probabilities are nonparametric, the procedure also becomes parameter-free.

Choosing the model with the smallest description length Σ is the means of balancing model complexity and goodness of fit^2^. We consider two candidate models throughout our experimental design: model *ℋ_a_,* where layers are true descriptors corresponding to the weighted edges of our deficit of interest in one layer and the connection matrix of the set of lesions in another layer, and a null model *ℋ_b_* where the edges describing deficit and the lesion connectivity effects are randomly interspersed across layers. The comparative magnitude of the description length of each model yields the following posterior odds ratio:

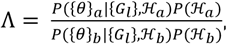

simplifying to

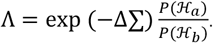

In this instance, *P*({*θ*}|{*G_l_*}*, ℋ*) is the posterior according to a given hypothesis *ℋ*, i.e., the true or null layered formulation. *P*(*ℋ*) is then the prior belief for hypothesis *ℋ*, and Δ Σ = Σ*_a_* − Σ*_b_* the difference in the model description length for these hypotheses.

### Semi-synthetic evaluation of stochastic block model lesion-deficit mapping

The application of stochastic block modelling (SBM) to lesion-deficit inference is underwritten by the validity of the underlying statistical framework. No purely empirical external validation is possible here because the functional anatomy of the neural substrate is definitionally unknown—it is what we are using lesion-deficit models to infer—and discriminative models of outcome need only identify the anatomical boundaries—jointly defined by lesion and neural effects—that determine outcomes. But we can establish a semi-synthetic validation framework, where an array of plausible, empirically-guided anatomical ground truths are hypothetically posited, and the fidelity of candidate models of real lesions used to retrieve them is explicitly quantified^4^. Here we derive ground truths from the large-scale meta-analytic repository NeuroQuery^5^, extracting functional maps of terms selected to span a broad range of cognitive domains, and exhibit widely distributed neuroanatomical patterns. The terms used were “action”, “aversion”, “language”, “mood”, “motor”, and “sensation”. Each map was re-sampled into the same 4mm^3^ isotropic space employed in the SBM pipeline, and thresholded at a conservative Z score of >=4. The connected components of each mask where identified, and clusters smaller than 27 voxels or sampled by fewer than 3 lesions were removed. Six sets of hypothetical continuous lesion-deficit relations were then created by summing over the intersection between each lesion segmentation and each corresponding ground truth, yielding a weighted “deficit score” for each patient and each functional domain.

We proceeded to evaluate these semi-synthetic lesion-deficit relations with a non-parametric Bayesian hierarchical weighted stochastic block model incorporating layered and attributed properties, implemented in graph-tool (https://graph-tool.skewed.de)^2, 3, 6–8^, exactly as in the main analysis. We began by fitting a null model, with the two kinds of edge weight—the deficit score and the lesion co-occurrence—randomly distributed across two layers. We then fitted a test model where each type of weight was consistently assigned to its own layer. Deficit weights were modelled as Gaussian; lesion co-occurrence weights as Poisson distributions. Having initialised a fit, we used simulated annealing to further optimise it, with 1000 iterations and a default inverse temperature of 1 to 10. We used model entropy to determine if the layered model fit was better than the null, indicating that the inferred community structure corresponded to the synthetic ground truth and lesion co-occurrence effects. To visualise the inferred communities, we backprojected the incident edge weights onto the brain, deriving the mean and 95% credible intervals. Bayesian model comparison based on minimum description length was used to determine if the layered models were more plausible than the null (Supplementary Figure 4). To show the relationship between inferred deficit and lesion co-occurrence edge weights, we extracted the posterior means whose 95% credible interval did not cross zero in each test model and plotted their correlation.

To compare the fidelity of our anatomical retrieval to that achievable with conventional lesion-deficit mapping, we performed standard mass-univariate voxel-wise lesion-deficit mapping (VLSM) of the same data implemented in SPM. The lesions were smoothed with a Gaussian kernel of 4mm full-width at half-maximum to facilitate voxel-wise spatial inference within a random Gaussian fields framework, and entered into a voxel-wise general linear model with the lesion intensity as the dependent variable and the deficit score as the independent variable. The resultant statistical maps were thresholded at p<0.05 family-wise error corrected. Models omitting the smoothing step yielded very similar maps (data not shown). The comparative fidelity of SBM vs VLSM models of the same data was then quantified by the difference in the Dice score for each inferred map relative to the ground truth. Note that the complex spatial structure of the ground truth maps employed here, chosen to provide the most robust test of the retrieval of distributed neural substrates, precludes interpretation of the absolute Dice score: the focus here is on the comparison with conventional topological models.

The SBM models were quantitatively superior to VLSM across the entire set (*p* = 0.028, Supplementary Figure 5). The qualitative results, visualised in the same figure, demonstrate the vulnerability of VLSM models to spatial biases driven by lesion morphology, and the ability of SBM models to resist them.

---

**Supplementary Figure 1.**
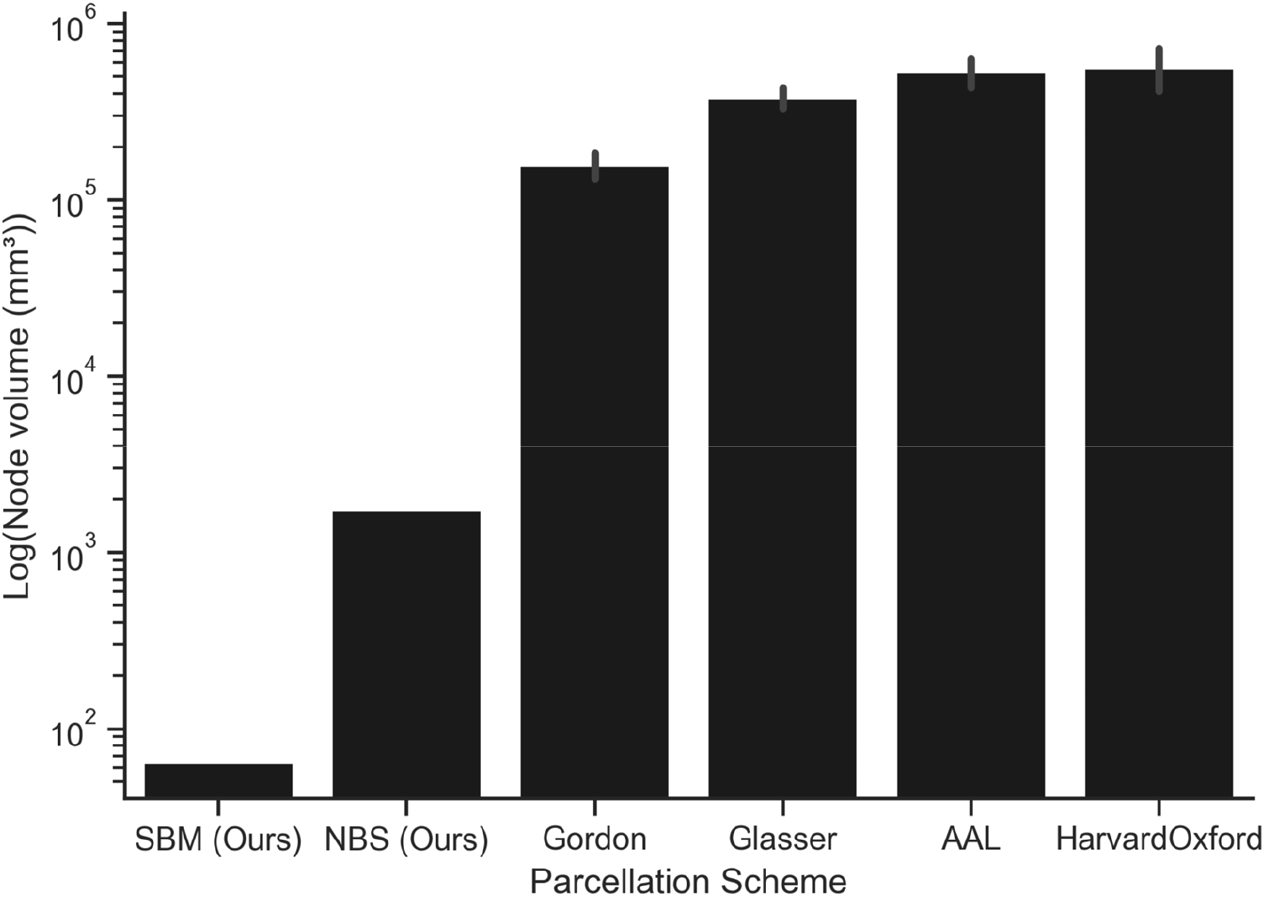
Bar plot of the mean spatial unit of analysis volumes of our stochastic block (SBM) and network based statistics (NBS) models compared with alternative regional parcellation schemes. Note our volumes are substantially smaller. The errors identify 95% confidence intervals. Note the ordinate is decimal log transformed.

**Supplementary Figure 2.**
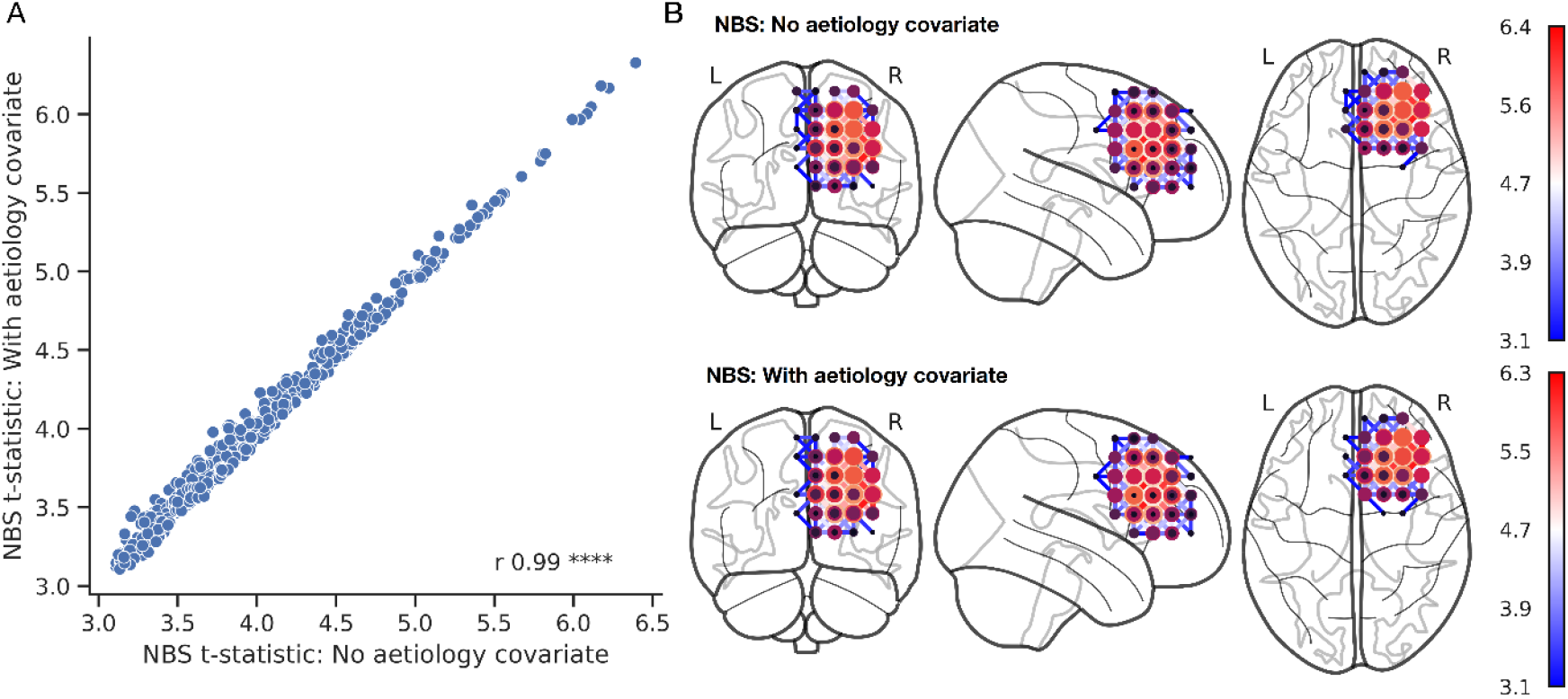
**A.** Scatter plot of significant edges derived from network-based statistics models including (ordinate) and excluding (abscissa) a lesion aetiology covariate. Note all points lie very close to the diagonal, indicating a minimal difference between the two. **B.** Visual connectome plots of the network-based statistics models excluding (top) and including (bottom) the aetiology covariate.

**Supplementary Figure 3.**
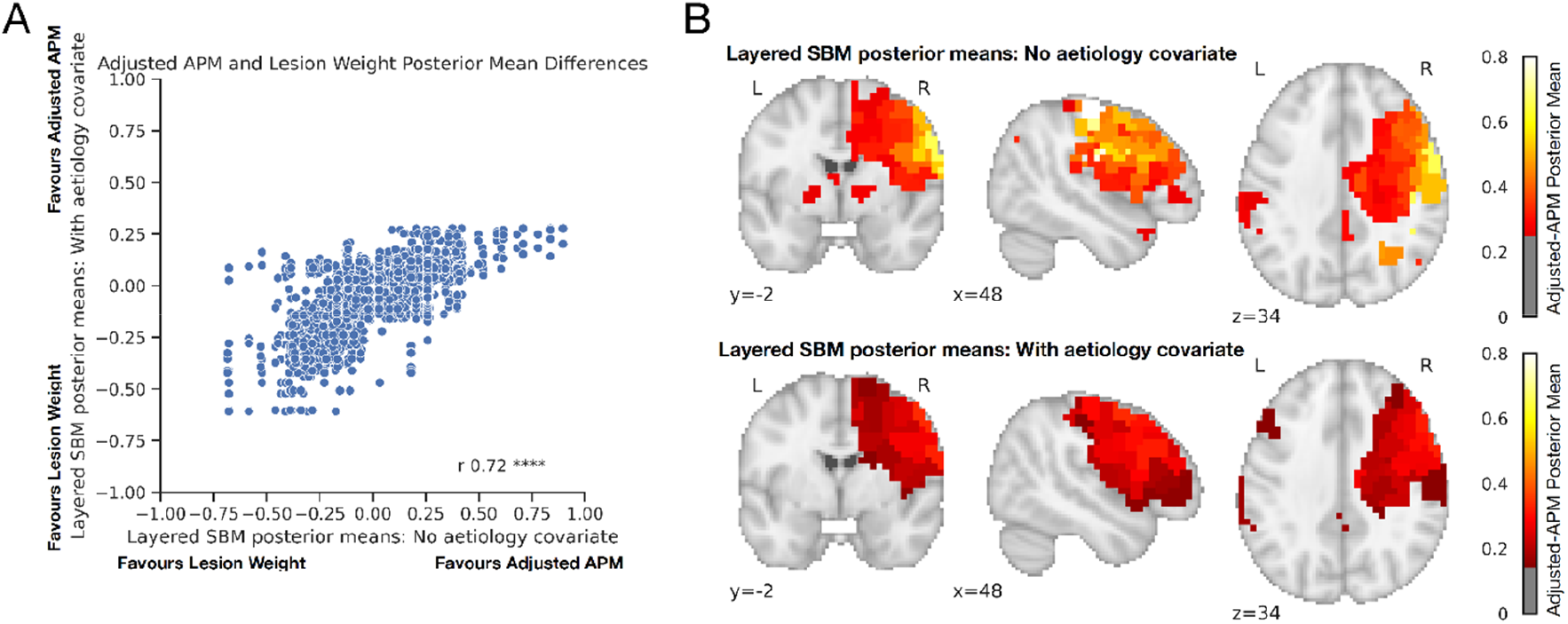
**A.** Scatter plot of the relation between posterior means of community blocks of stochastic block models with (ordinate) and without (abscissa) an aetiology layer. Note the former exhibits a narrower range of APM-related variation, suggesting weaker disentanglement from co-occurrence. **B.** Visualisation of the posterior means of the SBM without (top) and with (bottom) inclusion of aetiology in a separate layer. The description length of the more complex model was ∼2.4 times that of the simpler model (881118.22 vs 2133947.48 nats), a dramatic rise in description length for only a single additional feature, suggesting the former is overparameterised.

**Supplementary Figure 4.**
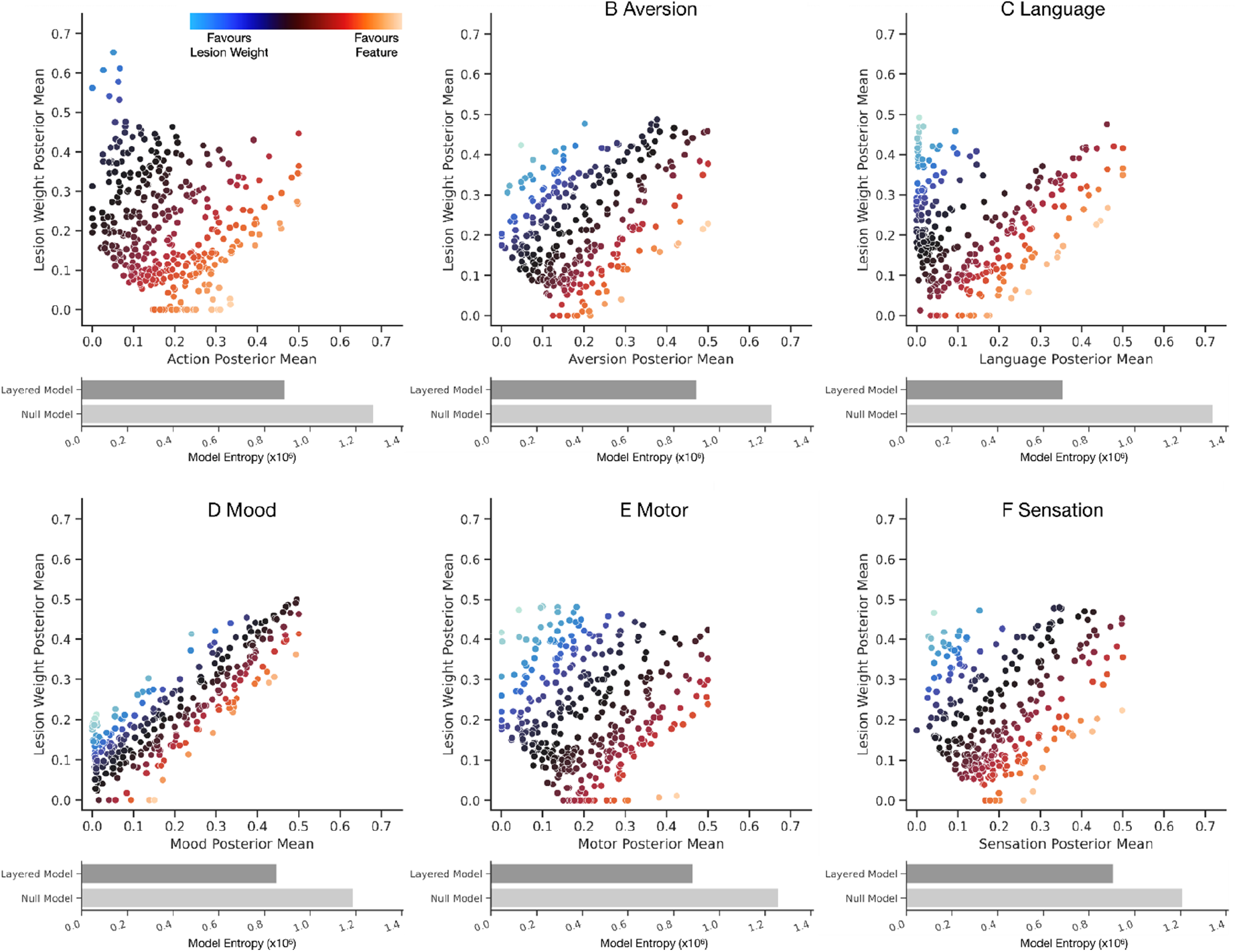
Scatter plots of the correlation between lesion co-occurrence (ordinate, red-yellow) and deficit (abscissa, blue-light blue) posterior means from the blocks of the test SBM models in each of the six ground truth experiments. Each individual point corresponds to a block at the l0 hierarchical layer. Below each plot is a bar chart of the corresponding test (dark grey) and null (light grey) model entropies in nats. Note the disentanglement of lesion co-occurrence and deficit effects, and lower entropies for the layered models across the set, indicating superiority. The entropy difference, x, between the layered and null formulation translates to a posterior odds ratio of e^x^ for the layered formulation over the non-layered alternative, as is further detailed in the supplementary methods.

**Supplementary Figure 5.**
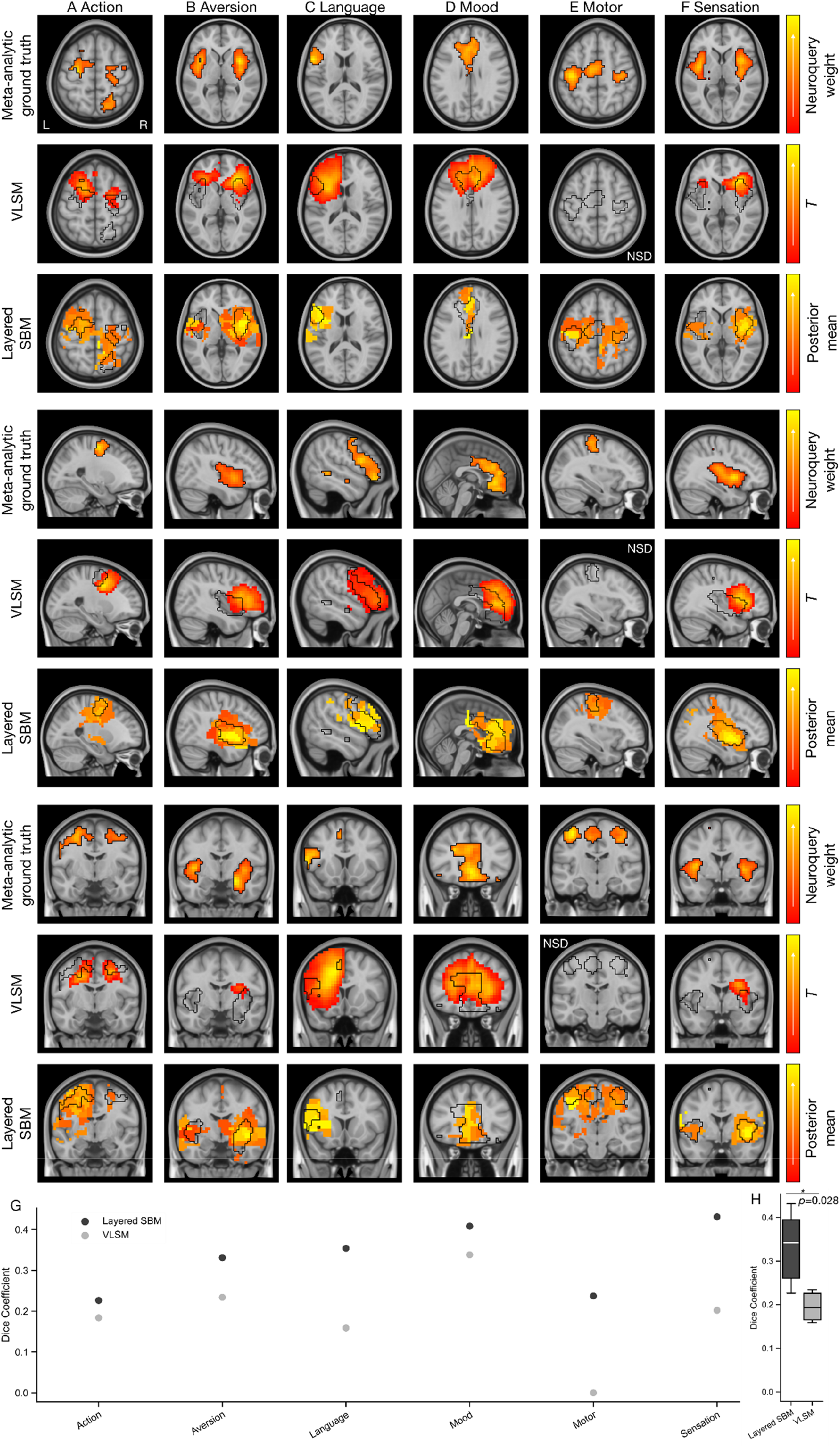
**A-F.** Triplanar view of the comparison between VLSM (rows 2, 5, and 8) and SBM (rows 3, 6, and 9) inference across the six domains, with the ground truth outlined in black over the source NeuroQuery weights (rows 1, 4 and 7). Note that VLSM retained no significant results for identifying the motor ground truth. **G.** Dice scores for SBM (black) and VLSM (grey) ground truth retrieval. **H.** Dice score boxplots across the set, with the p value for the difference.

